# Fine mapping of goat polledness variant in six Chinese native breeds

**DOI:** 10.1101/2021.12.20.473573

**Authors:** Yong Li, Tao Chen, Man-Man Yang, Hu Han, Dan jiang, Qiang Wei, Xing-Ju Zhang, Yan Ao, Qingfeng Zhang, Ze-Pu Miao, Ran Wang, Yuan-Lun Li, Sheng-Yu Chao, Lin Li, Ting-Ting Zhang, Ming Fang

## Abstract

**Background:** The genetic mechanism of goat polledness has been studied for decades, but identifying causative variants and functional genes remains challenging.

**Results:** Using a genome-wide association study (GWAS), we identified a significant striking locus for polledness in two different goat breeds. To reduce the linkage disequilibrium among variants for localizing causative variants in the finer region, we sequenced 79 goats from six Chinese native breeds (Jining Gray, Matou, Guizhou black, Yunnan black bone, Chaidamu, and Ujumqin) and identified 483.5 kb CNV (150,334,567-150,818,099) translocated into the previously identified 11.7 kb polled intersex syndrome region, which was consistent with previous research using intersex goat populations. Within the 483.5 kb CNV, a ~322 bp horn-specific element, similar to the superfamily of tRNA-derived families of SINEs, located at the first intron of the *ERG* gene was identified. The results of the GO enrichment analysis showed that the Horn-SINE element-associated genes were involved in both nervous system and head development. Finally, we used RNA sequencing to investigate gene expression profiles in the horn bud and skin tissues of horned and polled goats. We identified 1077 and 1222 differentially expressed genes between the horn bud and skin tissue in polled and horned goats, respectively. We also identified 367 differentially expressed genes in horn buds between polled and horned animals, and found that the two CNV-related genes, *ERG* and *FOXL2*, were upregulated in the horn bud of polled goats. Gene functional enrichment analysis demonstrated that the downregulated genes in the horn bud of polled goats were enriched in skeletal system development, whereas the upregulated genes were significantly overexpressed in muscle tissue development.

**Conclusions:** Broadly, this study describes a novel structural variant responsible for polledness/intersex traits and contributes to the discovery of molecular mechanisms underlying the development and regulation of the polledness trait.

## Background

Breeding of polled animals is an important goal for farming horned animals, such as cows, beef cattle, sheep, and goats, for animal welfare and economic reasons. There are several polled cattle and sheep breeds, such as Angus, Galloway, and hornless Dorset, in which the autosomal dominant polled allele has already been fixed^[1–3]^. However, the hornless goat breed has never been bred successfully, although goat polled markers have been extensively used in many countries^[4]^. In 1944, it was reported that goat polledness is closely linked with intersexuality. It is noteworthy to mention that intersexuality is a recessive trait, whereas polledness is a dominant trait^[5]^. The homozygous polled allele is accompanied by the generation of sex-reversing effect in XX individuals or mechanical obstruction of the epididymis for some XY individuals, resulting in breeding failure in hornless goats^[4]^.

The molecular mechanism responsible for development of polledness and intersexuality in goats has been studied extensively. Vaiman et al. found four microsatellite markers at 1q43a associated with the polled/intersex synchome (PIS) based on linkage analysis^[6, 7]^. Using a positional cloning approach, Pailhoux et al. identified an 11.7 kb deletion that triggered intersexuality and polledness by modulating the expression of *PISRT1* and *FOXL2*^[8]^ Gene knockout indicates that *FOXL2*, rather than *PISRT1*, is the causal gene responsible for the intersex of XX individuals^[9]^. It was also found that the expression of *PISRT1* and *FOXL2* was significantly increased in the horn buds of heterozygous or homozygous individuals without horns^[8]^. In addition, genome-wide association mapping of the polled locus revealed a strong signal on chromosome 1^[10]^. However, there is no direct evidence that *PISRT1* and *FOXL2* are responsible for the formation of polledness.

The formation of the horn is a very complex biological process that involves the differentiation and remodeling of various tissues, including the keratinization of the horn bud epidermis, ossification of the dermis and hypodermis of horn buds, and fusion with the skull^[11]^. The formation of horns is a successful evolutionary event. Recent studies have shown that pecorans with headgear-specific regulatory elements may play an important role in early cell remodeling during headgear development^[12]^. However, certain signaling pathways may interfere the process of horn development, which in turn leads to abnormal cellular remodeling, thus resulting in polledness. The causative genes underlying polledness have been identified for some species. A 1.8 kb insertion in the 3’ untranslated region (3’ UTR) of *RXFP2* has been identified to be associated with the horn/polledness of sheep^[13]^; *RXFP2* pseudogenization has been identified in Moschidae and Hydropotinae^[12]^, and a 212 bp insertion near gene *OLIG1* (~65 kb away) was found to produce hornless cattle^[14]^. Recently, two groups reported that a novel intersexuality-associated variant consisting of ~0.48 Mb duplicated fragment (including *ERG* and *KCNJ15*) downstream of the ~20 Mb PIS region was reversely inserted into the *PIS* locus of goat^[15, 16]^. However, the development of polledness trait in goats is very special because it is closely related to intersexuality, which increases the difficulties in the identification of hornless-related genes.

This study aims to employ sophisticated sequencing techniques to identify causative variants and to discover the genetic mechanisms underlying polledness trait development in goats.

## Materials and Methods

### Animal resources

Six native Chinese goat breeds were selected. All 381 Jining gray goats were sampled from Jining, Shandong Province, China, including polled (66) and horned (316) animals. A total of 735 Wushan white (WSW) goats were sampled from Wushan, Chongqing Province, China. Of these, 592 had horns and 143 were polled. Eleven Yunnan black bone (YNBB) goats (including four horned and seven polled goats) and eight Guizhou black (GZB) goats (including five horned and three polled goats) were sampled from southwest China. Ten polled Matou (MT) goats were sampled from Shangqiu, Henan Province, China. Eleven Ujumqin cashmere (UC) goats were sampled from Ujumqin, Inner Mongolia, China, including polled (3) and horned (8) animals. The sixth population was 30 horned Chaidamu cashmere (CDMC) goats sampled from Haixi, Qinghai Province, China. The sampling of each animal involved the collection of 1 ml of whole blood. All animal procedures were approved by the Life Ethics and Biological Safety Review Committee of BGI and were carried out following the approved guidelines.

### Genotyping and quality control

DNA was extracted from blood samples. The genomic DNA of each sample was digested with 1 μl Fast Digest TaqI (Fermentas; Thermo Scientific, Waltham, MA, USA) for 10 min at 65 °C in a reaction volume of 30 μl. For the ligation reaction, 1 μl of barcoded adapters (10 μM) was added to individual wells, along with T4 DNA ligase (Enzymatic) in a total volume of 40 μl. The ligation reaction was incubated for 1 h at 22 °C and heat-inactivated at 65 °C for 20 min. Twenty-four ligation products for different samples were pooled into a single tube, and 2 μl of chloroform was added to inactivate the restriction enzyme. The mixtures were centrifuged at 12000 rpm for 1 min, and the supernatant was transferred to a new tube. DNA fragments between 400-700 bp were selected on a 2% agarose gel (Amresco) and purified using a QIA quick Gel Extraction Kit (Fermentas; Thermo Scientific, Waltham, MA, USA). Samples were resuspended in 50 μl elution buffer and amplified by 10 cycles of PCR. The amplified library was purified using a QIA quick PCR Purification Kit (Fermentas; Thermo Scientific, Waltham, MA, USA), quantified on the Agilent2100 Bioanalyzer, and sequenced on an Illumina Hiseq2000 instrument with PE90 (Jining gray goat) or MGISEQ-500 sequencer with PE100 (WSW goat). The clean reads were mapped to the goat reference genome (ASM170441v1) (https://www.ncbi.nlm.nih.gov/genome/?term=capra_hircus) using BWA software (version 0.7.12); Samtools software (http://samtools.sourceforge.net/) was used to generate the consensus sequences for each goat and prepare input data for single nucleotide polymorphism (SNP) calling with realSFS (version 0.983) ( http://www.popgen.dk/angsd/index.php/RealSFS), based on the Bayesian estimation of site frequency at every site. Raw SNPs with sequencing depth greater than 2500 or less than 50, mostly resulting from repetitive regions or alignment errors, were removed for SNPs with extreme sequencing depth. An SNP was removed if its call rate was lower than 80%, its minor allele frequency (MAF) was less than 1%, or the proportion of its heterozygous genotypes was more than 60%; then SNPs were imputed with fastPhase (version 1.4) and Beagle software (version 5.1). The variants were filtered following the criteria: MAF >0.01, imputation information score > 0.9, and p-value of Hardy-Weinberg equilibrium (HWE) > 1e-6.

### Genome-wide association study (GWAS)

A total of 381 JNG and 735 WSW goats were collected, including 908 horned and 209 polled goats. Polled and horned animals were assigned as the cases and controls, respectively. The linear mixed model was used for the association study, *y* = *μ* + *Xb* + Zα + e, where *y* is the vector of phenotypes, μ denotes the overall mean, α represents the polygenic effect, and b is the estimator of fixed effects; e is a residual error, assumed to follow a normal distribution e~N (0,σ_e_), and X and Z are incidence matrices for b and α, respectively. The GWAS was conducted using the EMMAX software (http://csg.sph.umich.edu//kang/emmax/download/index.html). The significant threshold for association was set as 0.01 divided by the SNP number.

### Whole genome sequencing for fine mapping

Nine JNG, including four horned and five polled goats, were collected from Jining City, Shandong Province. The whole genome of each goat was sequenced using the BGISEQ-500 platform (PE100). The total sequencing amount for nine goats was 620.88 GB, and the sequencing reads were mapped onto the reference genome using BWA software.

In addition to JNG, five additional domestic goat breeds were collected (Additional file 1: **Table S1 and** Fig. S1). The genomic DNA of these sampled goats was extracted from blood or ear tissue and sequenced using the BGISEQ-500 or BGISEQ-2000 platform (PE100). The average sequencing coverage of CDMC, GZB, UC, JNG, YNBB, MT were 10.37×, 8.86×, 12.07×, 23.65×, 21.7×, and 10.87×, respectively. The raw reads were filtered with SOAPnuke1.5.0 (https://github.com/BGI-flexlab/SOAPnuke) and cleaned reads were mapped to the reference genome (GCA_001704415.1 ARS) using Bwa-0.7.12 (https://sourceforge.net/projects/bio-bwa/files/); the variants were imputed with STITCH R package, and filtered following the criteria: MAF>0.05 and HWE<10^-6^.

### Long-read whole genome sequencing

One polled goat was selected from nine JNG, and 37.22 Gb length was sequenced using the PacBio sequencing platform; the average read length was 12.94 kb. The sequencing reads were stored in BAM format and converted to FASTA by BAM2FastA the software smrtlink_4.0.0.190159 (https://www.pacb.com/support/software-downloads/). The transformed sequencing reads were then aligned to the goat reference genome using the ngmlr-0.2.7 software (https://github.com/philres/ngmlr). Samtools were used to transfer the Sam files to BAM files. Finally, the IGV software (https://igv.org/) was used to view the sequencing reads.

### Sanger sequencing

One PacBio sequencing read spanning the breakpoint was selected. The sequencing reads were aligned to chromosome 1 of the goat reference genome (accession number: NC_030808.1) using BLASTN and then manually spliced. Primers were designed to span the fusion breakpoint using the Primer3-py package (https://www.yeastgenome.org/primer3). A set of three PCR primers that amplified a series of specific bands was designed for genotyping horns and polled alleles. Primer and protocol information is shown in Additional file 1: Table S2. The composition of the reaction mixture for the PCR was as follows: 10 μl Premix Taq (Ex Taq Version 2.0 plus dye, TaKaRa, Japan), 0.3 μl each of forward and reverse primer (10μM), 1 μl template DNA (20ng/ul), and 8.4 μl ddH_2_O. PCR was performed using S1000 (BIORAD, USA) under the following cycling conditions: 2 min at 98 °C, followed by 29 cycles of denaturation, annealing, and extension at 98 °C, 30 sec at 58 °C, and 35 sec at 72 °C and a final extension for 2 min at 72 °C. The fragment length of PCR products was detected by 1.2% agarose gel electrophoresis using 5 μl of 100 bp DNA Ladder (TaKaRa, Japan) and gel stain (TransGen Biotech, China). A gel image of the different genotypes is shown in Fig. 4. The products were sequenced by Sanger sequencing and compared to the manually spliced reference sequence to validate the gene fusion. Sequence chromatograms were aligned and analyzed using the SnapGene2.3.2.

**Fig. 1.**
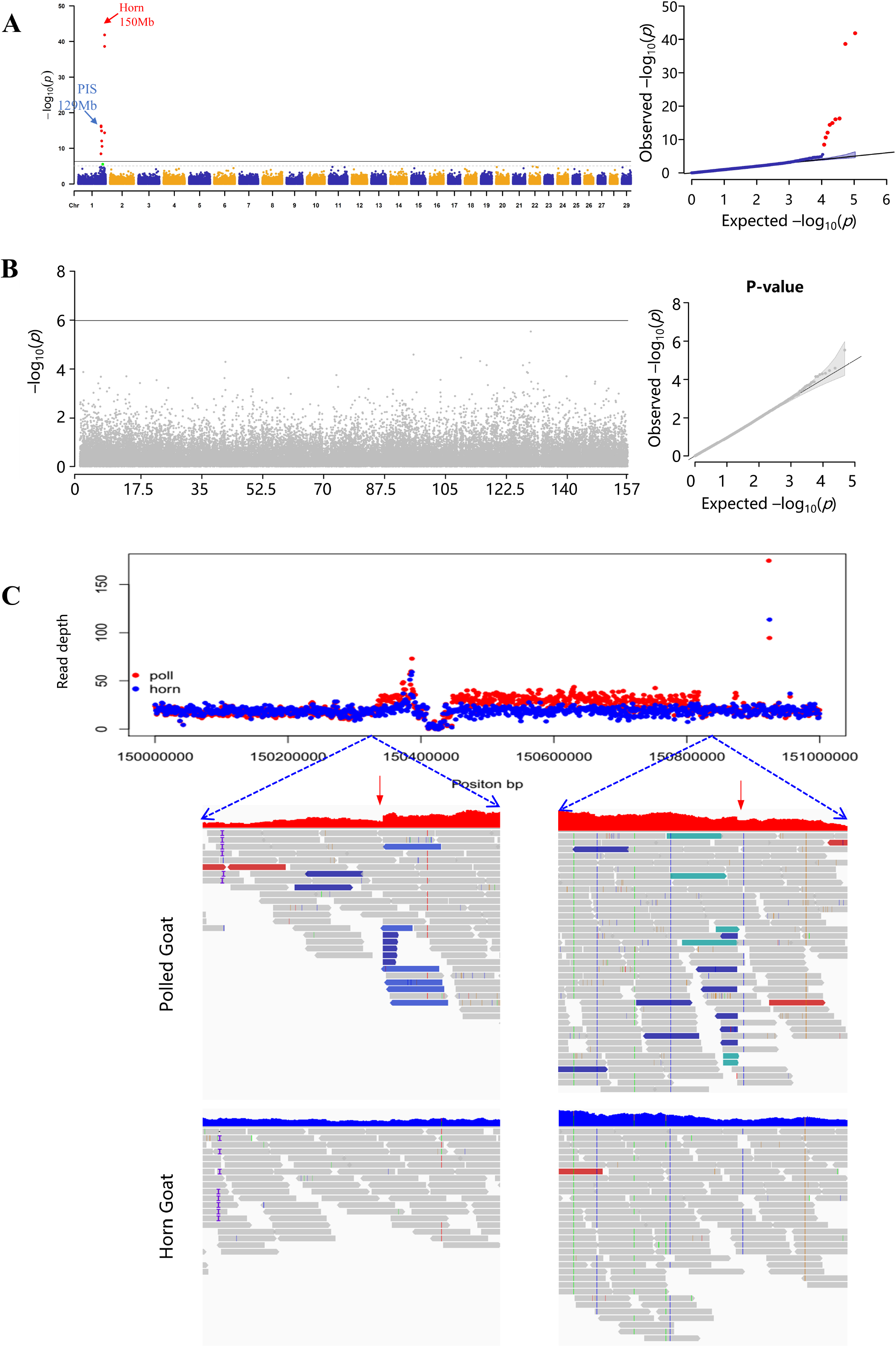
Genome-wide association and fine mapping study for identification of 483.5 kb CNV. (A) Genome-wide association study (GWAS) of polled/horn loci in Jining Gray (JNG) goat, the highest signal localizes at ~150Mb, and the next highest signal localizes at ~129Mb (PIS region). (B) The association study with 79 whole-genome resequenced goats from six breeds, 31 of polled and 48 horned, reveals that the signals at 150 Mb in the GWAS analysis are completely collapsed and no signals exceed the threshold. (C) The averaged read depths of five polled goats (red) and four horn goats (blue); the split reads events at breakpoints are shown below, it reveals that depth of reads of polled goats are strikingly higher than that of horn goats.

**Fig. 2.**
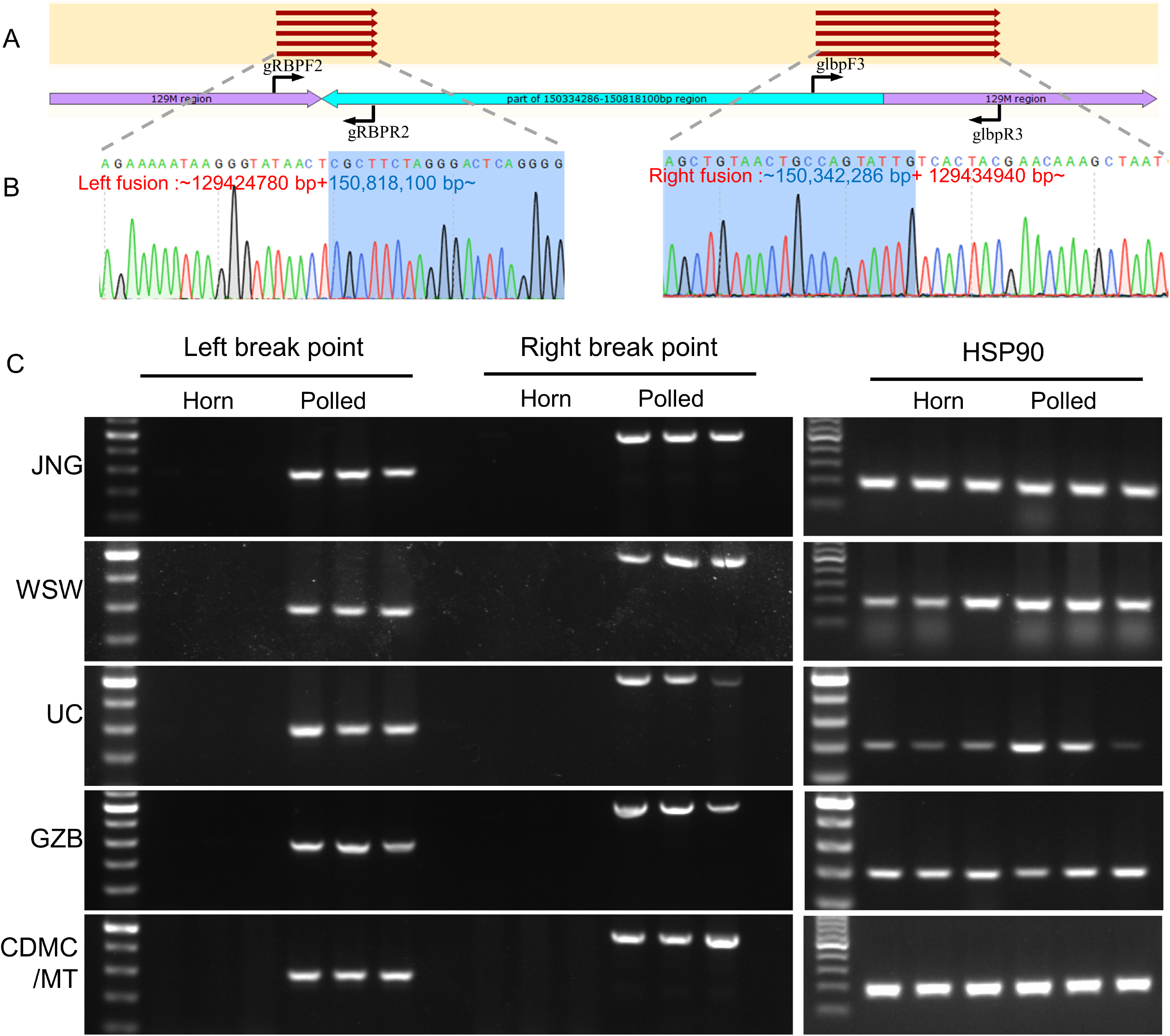
PCR validation of the translocation in six goat breeds. (A) Detection of PCR product for left and right breakpoints of horn and polled JNG goats. Horned goats with wild-type allele cannot be amplified, while polled goats have the duplication that allows amplification of the fragments at the junction. Hsp90 was used as a positive control. Arrows represent the position and direction of primers. (B) Sanger sequencing of the PCR products from gRBP2 and glbp3 bands. The results confirmed by Sanger sequencing were consistent with inferred sequence insertion patterns. The 483.5kb translocation is shaded in light blue. (C) Gel images of electrophoresed PCR products of the horn locus in horn and polled individuals of different goat breeds.

**Fig.3.**
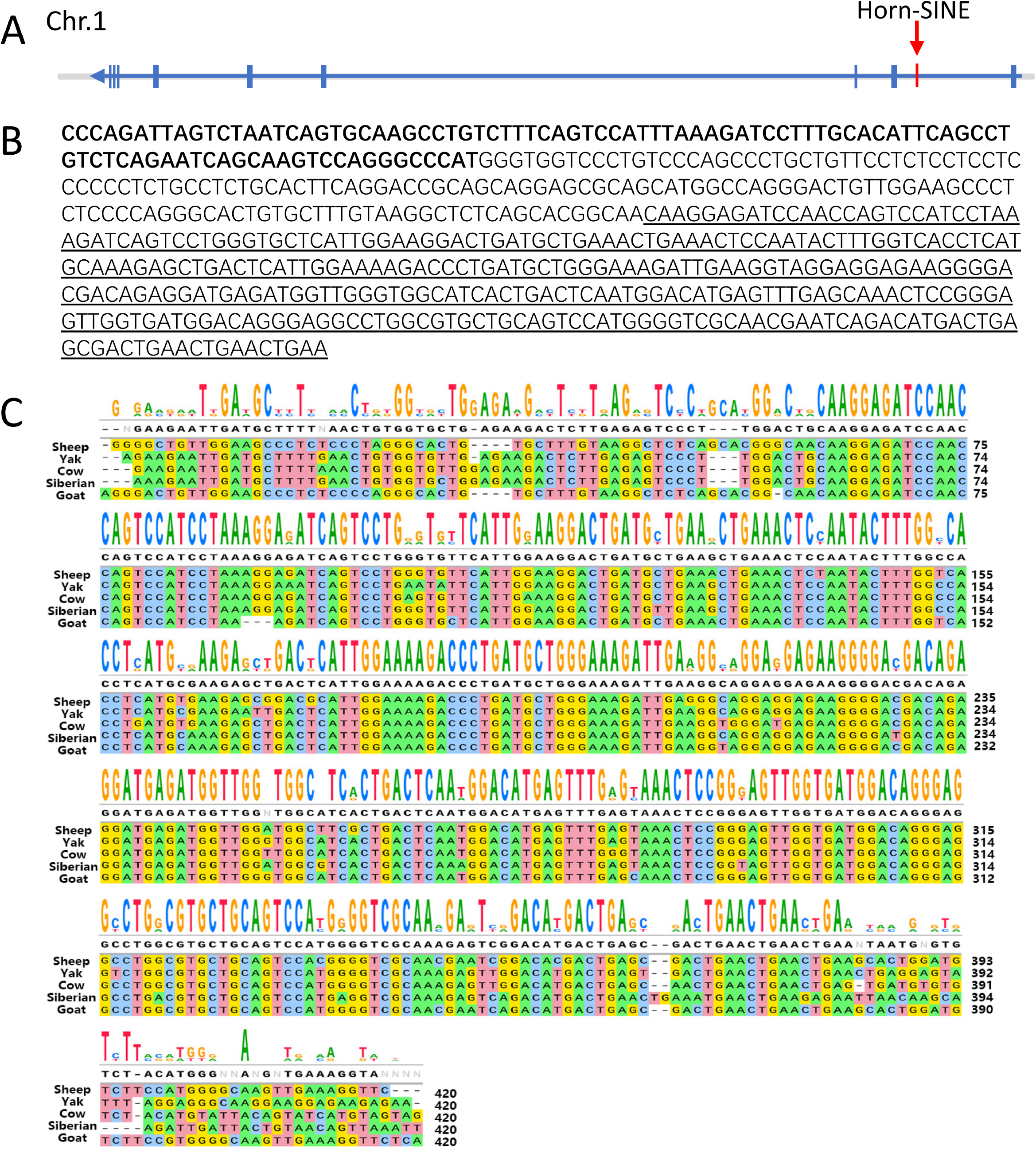
The nucleotide sequence and conservative analysis of the Horn-SINE element. (A) the Horn-SINE is located at the first intron of *ERG* gene. The schematic diagram indicates the gene structure of *ERG*. The red arrow indicates the Horn-SINE in our study. (B) The nucleotide sequence of the 5’ flanking region of the Polledness translocation t (1; 1). The PIS region sequence (129M) is in boldface. The Horn-SINE sequence is underlined. (C) Multiple sequence alignment of Horn-SINE sequence of five horned species (Goat, Sheep, Cow, Yak, and Siberian musk deer).

**Fig. 4.**
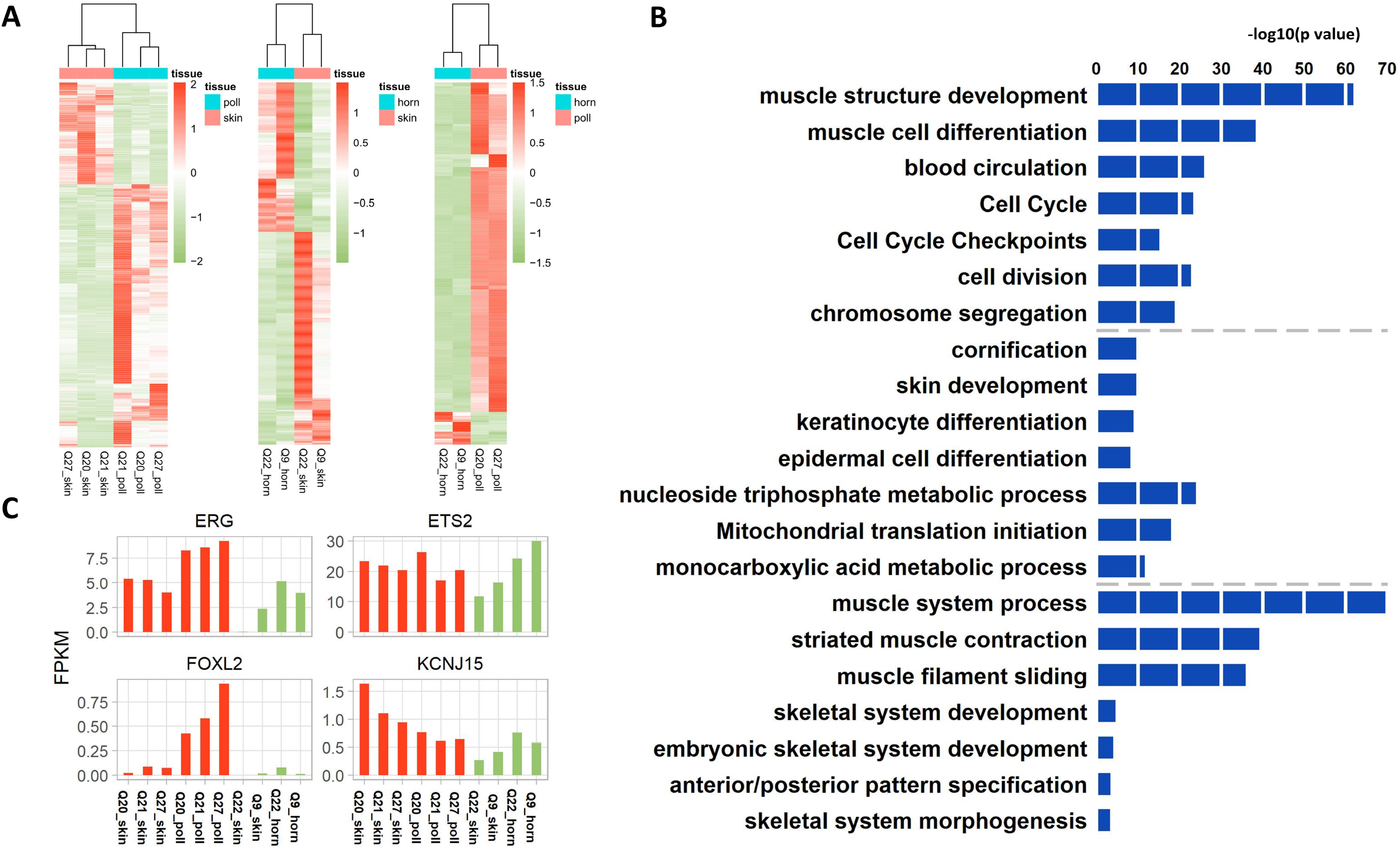
Abnormal Gene Expression in horn bud of polled goat at 4-day postnatal stage. (A) Heatmap illustration showing differentially expressed genes (DEGs) (p-adj <0.05) obtained by pairwise comparison between horn bud and skin of polled and horned goats, as well as between horn bud of horned and polled goats. (B) Gene ontology analysis of the differentially expressed genes in (A). Dashed lines separate different groups, GO enrichment of DEGs in up, middle, and bottom corresponding to left, middle, and right in (A); (C) The expression of *ERG, ETS2, FOXL2*, and *KCNJ15* in horn bud and skin tissue of horn (green) and polled (red) goats.

### SINE sequence analysis

Using BLASTN software, SINE-like sequences were aligned with the genomes of bison, cow, goat, antelope, yak, and sheep. Then we collected horn-specific SINE-like sequences that mapped to any of the selected species (E-value< 1e-5, identity ≥ 90%). After downloading the reference genome annotation file for the above species, gene annotation was performed in the 500 kb range around the horn-specific SINE-like sequence location. Finally, the annotated gene list was uploaded to Metascape (https://metascape.org/gp/index.html#/main/step1) for Gene Ontology (GO) and Kyoto Encyclopedia of Genes and Genomes (KEGG) enrichment analyses.

### RNA-sequencing

For RNA sequencing, we immediately collected ear specimens from five newborn Jining gray lambs and genotyped them by PCR using breakpoint primers. After that, two-horned (−/−) and three polled (inserted/−) lambs were euthanized by stunning at 4-day postpartum, and skin specimen and horn buds were collected and stored at −80 °C.

Total RNA was extracted from the skin and horn bud tissue of two horned and three polled goats using TRIzol reagent (Invitrogen, South San Francisco, CA, USA) following the manufacturer’s protocol. RNA integrity was evaluated using the Agilent 2100 Bioanalyzer (Agilent Technologies, Santa Clara, CA, USA), and 1.5 μg total RNA per sample was used as the input material for RNA sample preparation. The Illumina TruSeq RNA Sample Preparation kit (Illumina, San Diego, CA, USA) was used to construct transcriptomic libraries according to the manufacturer’s instructions [17]. Then, these libraries were sequenced on the Hiseq X10 platform at BGI (Shenzhen, China), and 150 bp paired-end reads were generated.

Sequence reads (paired-end, 150 bp) were trimmed using SOAPnuke (version 1.5.6). Then, the clean reads were aligned to the reference genome (GCA_001704415.1 ARS) using hisat2 (v2.1.0), and the number of reads mapped to the genome was counted using featureCounts. Based on the raw read counts, the R package DESeq2 was used to determine differentially expressed genes (DEGs), and reads with sum less than 2 was filtered out. Both corrected p-adj ≤ 0.05, and absolute log_2_ foldchange ≥ 1 value were set as the significant threshold for determining DEGs. Selected gene lists were subjected to GO term enrichment analysis using the Metascape database, and p-values were calculated based on the accumulative hypergeometric distribution.

### Availability of data and materials

The datasets generated and/or analyzed during the current study are available in the CNGB repository (CNSA: https://db.cngb.org/cnsa; accession number CNP0001896 and CNP0001914). The data that support the findings of this study are available from the corresponding authors upon reasonable request.

## Results

### Identification of candidate causal mutations for goat polledness in six Chinese native breeds

We collected 381 female Chinese Jining Gray (JNG) goats including 315 horned goats and 66 polled goats and further genotyped genome variants with double-digest restriction-site-associated DNA sequencing (ddRAD-seq), after which 107,012 SNPs were retained for the analysis. We then performed an association study using a mixed linear model that accounted for the polygenic effect using genomic kinships. Strikingly, a strong signal at ~150 Mb of chromosome 1 was identified (p =1.55e-42) (Fig. 1A). We then repeated the association with another goat breed of Wushan white (WSW) goats, including 592 horn goats and 143 polled goats genotyped with ddRAD sequencing. After quality control, 274,328 high quality SNPs were discovered. Evidently, the highest signal was localized at region 150Mb with p-values 1.79e-46 (Fig. S2), consistent with the results identified in the JNG goat population.

Next, we mapped the causative variant for goat polledness with whole-genome resequencing. We took advantage of multiple goat breeds to reduce the linkage equilibrium between variants. To do this, we collected 79 goats from six breeds, including 18 polled and 61 horned goats (Additional file 1: Fig. S1 and Table S1). All goats were sequenced with either Illumina Hiseq 2500 or the BGISEQ-500 platform with coverage of 8.86 to 23.65. After variant calling, filtration, and imputations, 14,978,368 variants were analyzed. Assuming that the polledness of different goat breeds was controlled by the same causative variant, we performed genome-wide association across breeds to identify polled loci using a linear model. However, the signal of 150 Mb on chromosome 1 that had been identified in JNG and WSW goat populations completely collapsed, and no variants reached a significant level (Fig. 1B). We hypothesized that the real causative variant did not exist in the variant set for the association study but instead linked with them in different linkage phases within each population, such that different directions of variants canceled the association signals. Furthermore, structural variants, such as copy number variation (CNV), which had not been genotyped with conventional bioinformatics pipelines, were used to identify causal variants for goat horn/polledness. By contrasting the read depths between four horned and five polled JNG goats, soon we discovered an obvious 483.5 kb CNV localized at ~150 Mb on chromosome 1 with a read depth ratio between the horn and polled groups ~2:3 (1:1.62) (Fig. 1C), suggesting that the horn goats had two copies of the 483.5 kb CNV, whereas the polled goats had three copies (heterozygous CNV). We then determined the breakpoints at single-base pair resolution using the split-read method. The results revealed that a 483.5 kb CNV locus on chromosome 1 (ASM170441v1, chr1:150,334,567-150,818,099), spanning the ETS transcription factor *ERG*, and overlapping potassium inwardly-rectifying channel subfamily J member 15 (*KCNJ15*). We additionally validated the 483.5 kb CNV with 70 goats from five breeds, including 23 polled and 47 horn goats. This CNV was present in 23 cases (100%) and 0 controls (0%) (Table1, Additional file 1: Fig. S3-21), suggesting the 483.5 kb CNV is most likely the causal variant for goat horn/polledness.

**Table 1.**
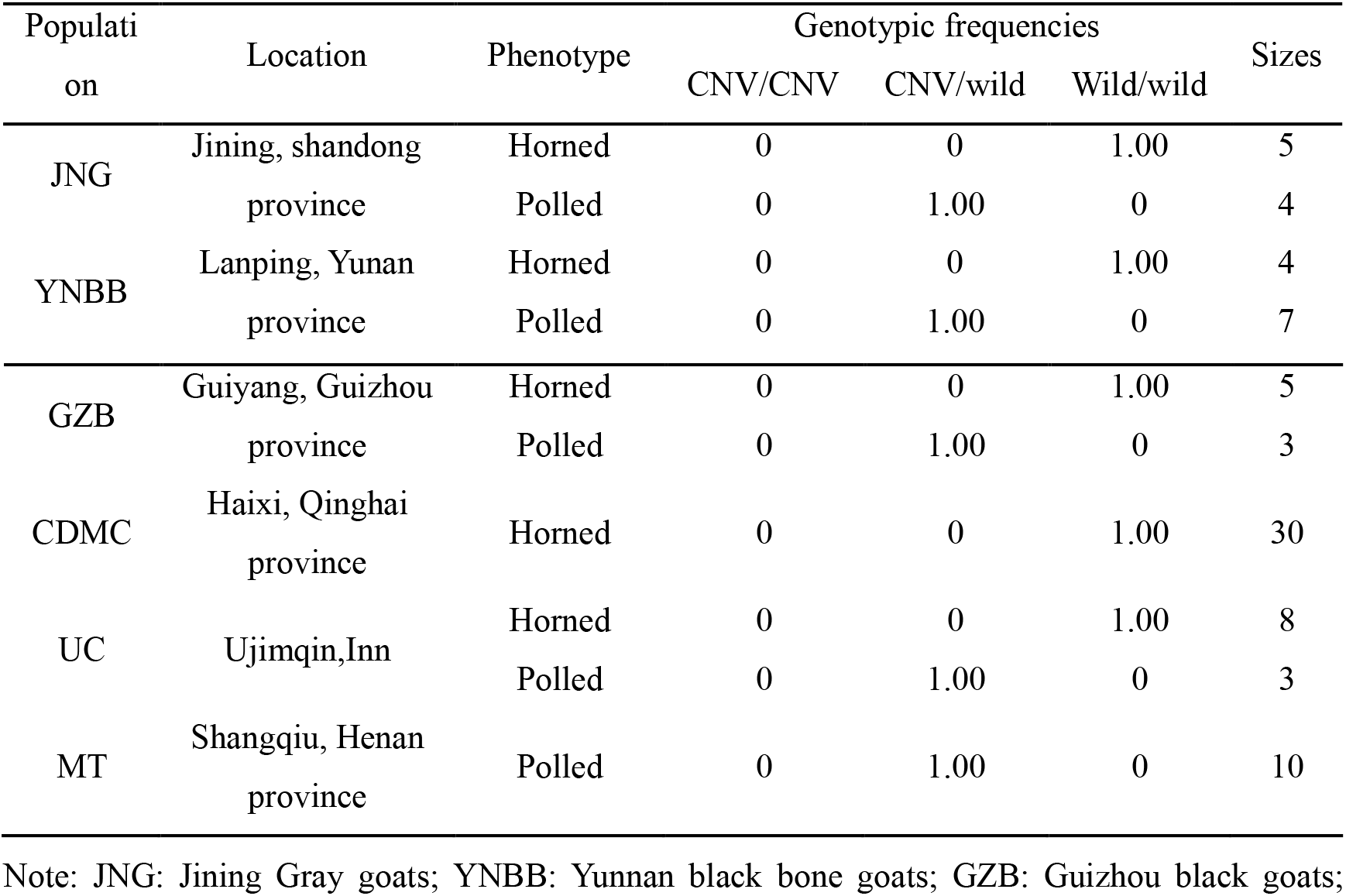
Frequencies of genotypes of 483.5kb CNV in six goat populations

Taking advantage of whole-genome resequencing, we re-identified the 11.7 kb deletion for PIS^[8]^ with single base-pair resolution (129,424,780–129,434,940bp) using the split-read method. When closely examining the breakpoints of the 11.7 kb deletion, some sequencing reads at the left/right outer edge of the 11.7 kb deletion were split and reversely mapped to the right/left inner edge of the ~150 Mb CNV (Fig. S22a, Fig. S23-S27). This suggests that the 483.5 kb CNV and 11.7 kb deletion are jointed, and the 483.5 kb deletion is reversely inserted into the 11.7 kb deletion to form a translocation (named Polledness translocation t (1; 1)). We then validated the Polledness translocation t (1; 1) using a long-read sequencing platform PacBio (average read length of ~12.94 kb). The results showed that this translocation was successfully re-identified (Fig. S22b), which strongly confirmed the reliability of the Polledness translocation t (1; 1). Finally, we applied the PCR-based method to validate the CNV in six JNG goats, including three polled and three horned goats. As expected, we obtained the target fragment for polled goats at either the left or right breakpoints (Fig. 2A, C), whereas no PCR products were amplified for horned goats (Fig. 2A, C). The PCR products of the polled gray goats were further validated by Sanger sequencing (Fig. 2B). We then validated the Polledness translocation t (1; 1) in other goat breeds, including randomly sampled 6 WSW (three polled and three horn goats), six UC (three polled and three horn goats), six GZB (three polled and three horn goats), three CDMCs (all horn goats), and three MT (all polled goats). The results showed that the target fragment of all polled individuals was amplified at either the left or right breakpoints, whereas no PCR products of horned individuals were amplified (Fig. 2C).

### Conservative analysis of the sequence of the Polledness translocation t (1; 1)

Analysis of sequence features flanking the breakpoints of the 483.5 kb CNV revealed a 322 bp conserved repeat sequence in the first intron of the *ERG* gene (150,817,512-150,817,809) (Fig. 3A, B). BLAST analysis showed that this sequence was a type of SINE sequence, similar to a tRNA pseudogene coupled to an A element (Bov-tA) (Additional file 1: Fig. S28), which has not been reported in the literature yet ^[12]^. The horned goats had two copies of the SINE sequence, whereas the polled goats had three copies (heterozygous) or four copies (homozygous). We then aligned the SINE sequence to the genomes of several horned animals such as cows, goats, sheep, chiru, and yak, and found that it is highly conserved in these horned species (Fig.3C).

Interestingly, this SINE sequence (named Horn-SINE) can also be identified in other regions of the horned species’ genome, with a BLAST identity score of more than 91%. Subsequently, we extracted the Horn-SINE overlapping genes in the genomes of goats, cows, sheep, chiru, and yak, corresponding to 195, 177, 321, 205, 148, and 76 genes, respectively (Additional file 1: Fig. S29; Additional file 2: s1). They were significantly enriched in the signaling pathway regulating the RAC1 GTPase cycle (adjusted P-value=10^-8.35^) and nervous system development (adjusted P-value=10^-6.26^) (Additional file 1: Fig. S29; Additional File 2: s4). The RAC1 GTPase cycle plays an essential role in osteoclasts by regulating actin dynamics, and nervous system development is also associated with horn growth^[18, 19]^. In addition, we extracted the genes in the 50 kb and 250 kb regions on both sides of the SINE-like element and analyzed the gene function category and enrichment (Additional file 2, s2, s3). The results showed that nervous system development, small GTPase mediated signal transduction, human cancer, and signaling by receptor tyrosine kinases were significantly enriched (Additional file 1: Fig. S30-31; Additional file 2: s5 and s6). To further elucidate the relationship between Horn-SINE element-related genes and horn formation-related genes, we downloaded the horn-specific expression genes of goat and sheep (data from PRJNA438286; Additional file 2: S7) and accessed DEGs between horned and polled bovines (data from Wiedemar^[20]^; Additional file 2: S8) from previously published transcriptome data^[12, 20]^. The horn-specifically expressed genes were enriched in signaling by receptor tyrosine kinases and head development (Additional file 1: Fig. S32; Additional file 2: S9), while the bovine DEGs were enriched in both nervous system and head development (Additional file 1: Fig. S33; Additional file 2: S10). Taken together, pathways regulating the nervous system and head development were enriched in almost all groups (Table. 2), demonstrating the reliability of our results.

**Table 2.** Enrichment analysis of SINE-like associated genes in horned animals

| Pathway | Overlap<br>ped | 100k<br>flanking | 500k<br>flanking | Horn-specific<br>genes | DEGs in<br>horned<br>cattle |
| --- | --- | --- | --- | --- | --- |
| Nervous system development | √ | √ | √ | √ | √ |
| RAC1 GTPase cycle | √ |  |  |  |  |
| Human cancer |  | √ | √ | √ |  |
| Signaling by receptor tyrosine<br>kinases |  | √ | √ | √ |  |
| Head development |  | √ | √ | √ | √ |
| Small GTPase mediated signal<br>transduction |  | √ |  |  |  |
| Skeletal system development |  |  | √ |  | √ |

**Notes:** overlapped: element located in the gene. 100k flanking: 50 kb flanking each side of the element; 500k flanking: 250 kb flanking each side of the element; Horn-specific genes: genes expressed explicitly in horn bud of goat and sheep; DEGs in horned cattle: genes differentially
expressed in horn buds between horned and polled cattle. **Abbreviations:** DEGs, differentially expressed genes.

### Gene expression studies reveal novel horn development-specific candidates

Initially, we investigated the DEGs at the horn bud between horned and polled goats and between facial skin tissues and horn bud. Five goats (three polled and two horned) were sacrificed at day 4 after birth and the horn buds and facial skin tissues were harvested for isolation of the total RNA, and the ten mRNA samples were sequenced to investigate the possible effects of the polled mutation on gene expression. Finally, 1077 DEGs between horn bud and skin tissue in polled goats were identified, among which 776 upregulated genes in horn bud tissues of polled goats were enriched in skeletal muscle tissue development and blood circulation. There were 1222 DEGs between horn bud and skin tissue in horned goats, and 504 upregulated genes were enriched in cornification and skin development (−log_10_(p-value) =9.86) (Fig. 4A, B and Additional file 3, S1-S2 and Additional file 4: S1-S4). In addition, 367 DEGs in horn buds between horned and polled goats were identified, which were enriched in muscle structure development (-log_10_(p-value) =63.00) and skeletal system development pathways (-log_10_(p-value) =4.59) (Fig. 4A, B and Additional file 3: Table S3 and Additional file 4: Table S5-S6). We next investigated the alterations in the expression of key genes near or on the Polledness translocation t (1; 1), including *KCNJ15, ERG, FOXL2*, and *ETS2*, among which *FOXL2* has been identified as a causal factor for PIS^[9]^. For genes in or near the CNV and 11.7kb deletion region, *ERG* and *FOXL2* were upregulated in horn bud tissues compared to skin tissue of polled goats, whereas *KCNJ15* and *ETS2* were not differentially expressed (Fig. 4C). Furthermore, we found that *ERG* and *FOXL2* were upregulated in horn bud tissues of polled goats compared to the horn bud tissues of horned goats (Fig. 4C). We also found that *ERG-*targeted genes, like *IGF2* and *IGFBP1*^[21]^ were upregulated, while some interactive genes, such as homeobox genes, *Hoxc6, 8, 9, 5*, and *Hoxb7, 6, 3*, as well as *Hoxa9, 7*^[22–24]^ were downregulated in horn bud tissues of polled goats (Additional file 3: Table S1). In addition, *FOXL2-*regulated genes, such as some collagen genes^[25]^, were upregulated in the horn bud tissues of polled goats too (Additional file 3: Table S1).

## Discussion

In this study, we found that for all the polled goats regardless of goat breeds, the 483.5kb CNV was inserted into an 11.7 kb deletion to form the Polledness translocation t (1; 1), but this translocation did not occur in any of the horned goats. Our GWAS and fine-mapping results were consistent with the two recent research results from Switzerland and China that used long-read whole-genome sequencing of two genetically female goats (a PIS-affected and a horned control) and whole-genome selective sweep of intersex goats from China, respectively^[15, 16]^. Although the results of the three groups are consistent, our study is different from the other two studies. We took advantage of multiple breeds to fine map causative mutations. We first used GWAS to identify a region using natural goat populations (381 JNG goats and 735 WSW goats), and used multiple goat breeds to exclude all SNP and INDEL variants and then used resequencing to identify structural variants. In contrast, the other two research groups used long-read whole-genome sequencing or selective sweep analysis to identify structural variants in intersex and non-intersex goat populations^[15, 16]^. Due to the close linkage of polledness and intersex, research based on different ideas could obtain similar results. This indicates the reliability of our results. In addition, we also verified the 483.5 kb CNV in more breeds of goats, indicating that goat hornless traits have a consistent genetic mechanism. Finally, the previous two articles only found more complex mutations in the PIS region and did not elaborate the molecular mechanisms of polledness.

PIS is a unique phenomenon in goats. There is no such phenomenon in other horned animals, indicating that gonad development and horn development are not necessarily related. The development of PIS is related to the complex structural variation in chromosome 1 of goats. Previous results showed that the deletion of 11.7 kb activated the expression of *FOXL2* in gonadal tissue, which in turn led to the occurrence of sexual reversal^[8]^. The gene knockout results confirmed that the intersex causative gene was *FOXL2*^[9]^. However, how the 11.7 kb deletion regulates the formation of polledness is still unknown. The 483.5 kb CNV is translocated into the PIS region, providing novel insights into the genetic mechanism of goat polledness.

Sequence conservation analysis and functional element identification showed that a SINE sequence was located in the first intron of the *ERG* gene near the left breakpoint, which is specific to horned animals. This Horn-SINE sequence is similar to a tRNA pseudogene coupled to an A element (Bov-tA), which belongs to the superfamily of tRNA-derived families of SINEs^[26, 27]^. These Bov-tA were established after the divergence leading to the establishment of Suidae and Bovidae families, and these SINE insertions may be informative for phylogenetic reconstructions of ruminants^[28]^. In addition, Horn-SINE also has multiple homologous copies in other genomic regions of horned species. These results indicate that Horn-SINE may have important biological functions in horn formation. SINEs are a class of retrotransposons transcribed by RNA polymerase III, which do not encode proteins. SINEs can function as cis-or trans-regulatory RNA elements that regulate gene expression from a distance as a tissue-specific enhancer^[29]^. Enrichment analysis of Horn-SINE-associated genes revealed that signal pathways related to neural development, RAC1 GTPase cycle, head development, signaling by receptor tyrosine kinases, and human cancer were enriched, which is consistent with other studies^[12]^. Further, we analyzed the goat and sheep horn-specific expression genes^[12]^ and the DEGs between cattle horned and polled^[23]^ using published data. The functional categories of these genes were highly consistent with the functional categories of SINE-related genes. These results suggest that the Horn-SINE is not an evolutionary trace formed by random insertion and is more likely to play a role in regulating horn development. This function may be achieved by regulating the expression of its surrounding genes; when translocation occurs, the 482.5kb CNV carrying SINE-like element results in abnormal expression of the causative gene in horn tissues, and this aspect is worthy of verification in subsequent experiments. In addition, we obtained only limited published transcriptome data and could not obtain transcriptome data at different developmental stages. In the future, we will analyze the expression patterns of Horn-SINE-related genes based on the comparative transcriptome data of horned and polled individuals at different developmental stages to further confirm the biological functions of Horn-SINE.

It is well known that CNV not only affects the expression of related genes, but also affects the expression of genes located near the rearrangements at distances of up to several hundred kilobases ^[30]^. This translocated and adjacent region contains three genes, *ERG, KCNJ15*, and *FOXL2*, of which *ERG* does not contain the first exon. Our RNA-seq results showed that *ERG* and *FOXL2* were upregulated in the horned tissues of hornless goats, and there was no difference in skin tissues between polled and horned goats, whereas *KCNJ15* and *ETS2* were not differentially expressed in all tissues. Previous studies have also shown that *FOXL2* is absent in the gonadal tissue but upregulated in the horn bud tissue, and there may be a gonadal-specific regulatory element in the 11.7kb deletion region, which can inhibit the expression of *FOXL2* in gonadal tissue through its secondary structure^[8]^. The described breakpoint in the region of Chromosome 1 at 129Mb is in the *FOXL2* topologically associating domain (TAD) when compared to the corresponding human genome region^[31]^. In contrast, the duplicated genomic segment contains the *KCNJ* gene and parts of the *ERG* gene, as well as parts of the respective TADs. When the duplication is inserted into the breakpoint, it can be assumed that a fusion TAD (neo-TAD) is formed, consisting of one part of the duplication and the rest of the *FOXL2* TAD. This was confirmed by one recent study, which showed that the inserted duplication changed the original spatial structure of goat *CHI1* and changed the appearance of several specific loop structures in the adjacent ~20kb downstream region of *FOXL2*^[16]^. Due to the inversion, *KCNJ* is placed on the other side of the boundary and isolated; thus, *KCNJ15* was not differentially expressed in all tissues between polled and horned goats. Therefore, it must be the “rest” of the *ERG* gene that is of functional importance. The *ERG* gene is a member of the erythroblast transformation-specific (ETS) family of transcription factors, which is expressed during the earliest events of skeletal formation and cooperates with *TGF-β* to regulate the differentiation of the sclerotome^[32]^. In humans, the *ERG* oncogene is frequently overexpressed due to chromosomal translocations, resulting in different fusion gene products^[33–35]^. Dysregulation of *ERG* can result in abnormal development of the chicken limbs^[36]^. We found that the two most abundant transcripts were transcribed from the 4th exon of *ERG* using intragenic promoters or regulatory elements in CNV (Additional file 3: Table S4). This fusion region also contains a horn-specific regulatory element that leads to the misexpression of *FOXL2* in the horn buds of polled goats. *FOXL2*, encoding a forehead transcription factor, plays a role in ovarian, skeletal, and muscle development^[37, 38]^. Previous reports revealed that *FOXL2* was overexpressed in the horn tissues of some other headgear animals, such as deer and cattle^[12, 20]^.

Horn development is the result of the differentiation and remodeling of various tissues, including the ossification of hypodermal tissue and keratinization of the horn bud epidermis^[1]^. In horned cattle, ossification of the developing horn occurs one month after birth, while it is suppressed in hornless cattle^[39]^. In our RNA-seq study, the results of GO or KEGG pathway analysis showed that most of the key genes involved in skeletal muscle development, like *PAX7, MYOD, MFY5, MYOG*, and *MIR133A*, were upregulated, while genes related to bone and nerve cell development, such as *HOXA9, HOXC9, HOXC5, PAPPA2*, and *OTOP3*, were downregulated in the horn bud of polled goats. The enrichment of these DEGs was associated with the abnormal development of goat horn. We found that upregulated expression of *ERG* and *FOXL2* lead to the upregulated expression of targeted genes, such as *IGF2*, *IGFBP1*, and collagen genes, which promoted the differentiation of skeletal muscle cells and inhibited the differentiation of bones and keratinocytes. Therefore, we assume a proposed model of hornless trait formation involving ERG- and FOXL2-related signaling pathway mediated by the Polledness translocation t (1; 1) (Fig. 5).

**Fig. 5.**
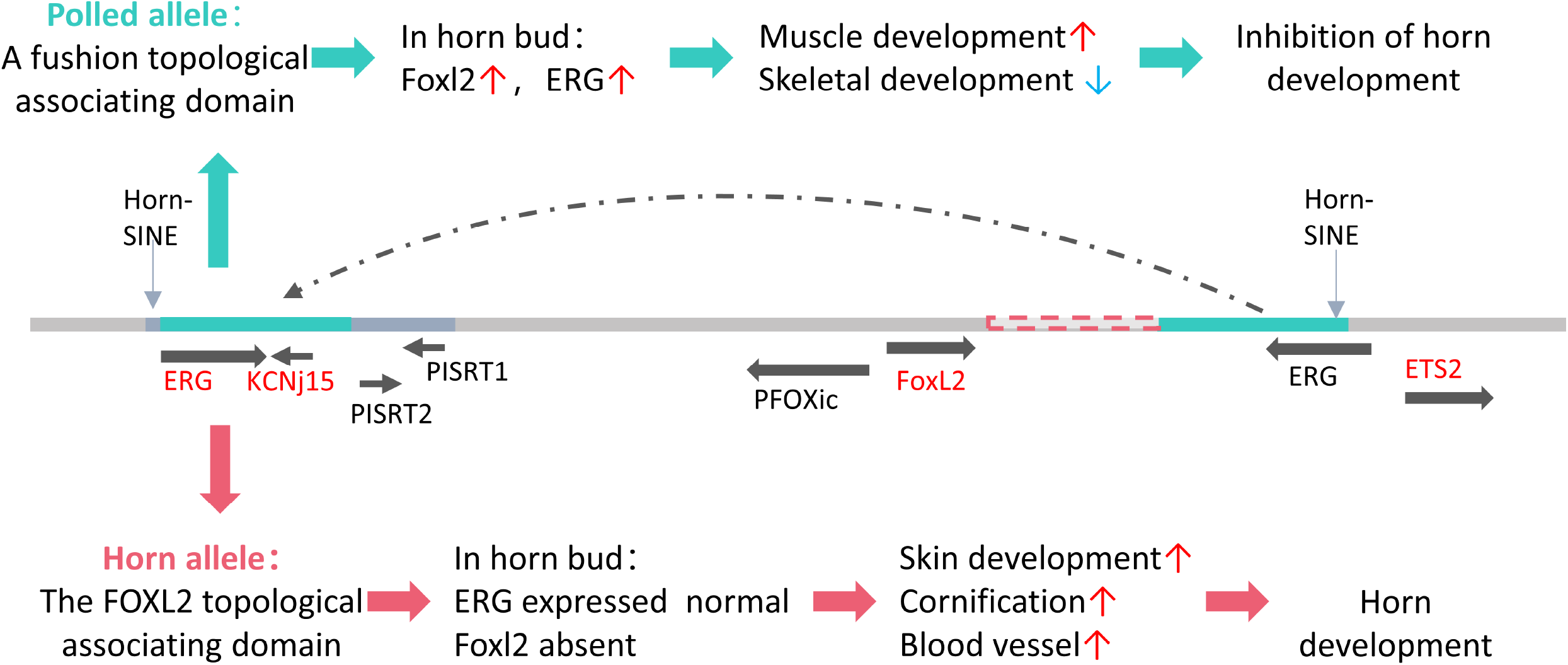
Proposed model of development of hornless trait involving *ERG-* and *FOXL2*-related signaling mediated by the Polledness translocation t (1; 1). We show here that the PIS deleted region is in the *FOXL2* topologically associating domain (TAD), which inhibits neighboring genes expression including *FOXL2* and *PIRSTs* et al., while *ERG* is expressed normally at its location far away from the TAD region in horned goats. In polled goat, the duplication is inserted into the breakpoint, and a fusion TAD (neo-TAD) would be formed consisting of one part of the duplication and the rest of the *FOXL2* TAD. This neo-TAD may upregulate *ERG* and *FOXL2* in the horn buds of polled goats. The expression change of two genes may repress expression of some horn development-related genes and activate expression of other tissue-related genes, like those involved in skeletal development and muscle development, which inhibits horn development. Together, these results introduce a novel genetic mechanism required for establishing the proper conditions for development of polledness trait.

## Conclusions

Our study provided evidence for a hornlessness-related 483.5kb CNV translocated into the previously identified 11.7kb PIS region, explaining why goat polledness is closely linked with intersexuality. We identified a horn-specific SINE-like element in the CNV region, and the enrichment analysis of the Horn-SINE-related genes revealed that the signaling pathways that were enriched are related to horn formation. We demonstrated a polledness formation model, in which horn-specific upregulation of two CNV-related genes, *ERG* and *FOXL2*, might repress the expression of some horn development-related genes and activate the expression of genes involved in skeletal development, muscle development, among others, which results in horn development dysregulation. It is imperative that we provide a novel genetic mechanism for the classical Mendelian trait found in the last century.

## Ethics declarations

This study was approved by the institutional review board of BGI (NO. FT 18041).

## Consent for publication

Not applicable.

## Competing interests

The authors declare that they have no competing interests.

## Funding

This work was supported by the Science and Technology Innovation Strategy Projects of Guangdong Province (2019B020203002), National Natural Science Foundation of China Grant (31872560 and 31672399); Key Research and Development Project of Tianjin (20YFZCSN00720)

## Authors’ contributions

L.Y conceived, designed, and supervised the study with F.M., C.T., H. H., F. M., L.Y., M.Z.P performed the informatics analysis of the sequencing data. Y.M.M., W.R., Z.Q.F., L.Y.L., C.S.Y., Z.X.J., L.L obtained goat material and DNA for resequencing and genome sequencing. Z.X.J., W.R., L.L., Z.T.T obtained goat tissues and RNA for RAN-sequencing. C.T analyzed the transcriptome. L.Y., F.M., H.H., C.T. and Y.M.M are the major contributors in writing the manuscript. All authors read, revised, and approved the final manuscript.

## Acknowledgments

We thank Hu Yeyong from Henan yudong animal husbandry co. LTD and Bayaertu & Yong quan from Dongwuzhumuqin Banner Aogali Animal Husbandry Co., Ltd provide MT goat and UC materials, respectively.

